# Targeting GDF15 as a potential therapy for low back pain originating from endplate

**DOI:** 10.1101/2023.02.11.528124

**Authors:** Xiaoqun Li, Jinhui Wu, Qingjie Kong, Miao Hu, Ziheng Wei, Heng Jiang, Zheng Zhang, Runze Gao, Xuhui Zhou, Jun Ma

**Author notes:** Corresponding author and to whom requests for offprints should be sent: Xuhui Zhou and Jun Ma, Address: Department of Orthopedics, Shanghai General Hospital, Shanghai Jiao Tong University School of Medicine, No.85 Wujin Road, Shanghai, P.R.China. Lixiaoqun, Jinhui Wu, Qingjie Kong and Miao Hu contributed equally to this work.

## Abstract

The overactivation of osteoclasts in the endplate is one of the most important causes of low back pain (LBP) originating from endplate. Transforming growth factor-β family has been demonstrated to play an important role during osteoclast differentiation. GDF15 was reported to participate in several pathological states. In this study, we reported that the lumbar spine instability (LSI) induced the overactivation of osteoclasts and CD31^hi^Emcn^hi^ endothelial vessels in the bony endplate. In addition, the expression of GDF15 in human disc samples and mice models were also increased. GDF15 could promote the fusion of preosteoclasts via Rac1/Cdc42/PAK/Cofilin axis, and facilitate the angiogenesis via the secretion of PDGF-BB. Furthermore, we proved that the GDF15 inhibitor, CTL-002 could reduce the expression of GDF15 in the endplate and alleviate the overactivation of osteoclasts and CD31^hi^Emcn^hi^ endothelial vessels induced by LSI in vivo. In conclusion, we demonstrated that GDF15 could regulates the fusion of preosteoclasts and targeting GDF15 in the endplate served as a novel anabolic therapy for low back pain treatment.

## Introduction

Low back pain (LBP) is a serious social health and clinical problem which impact over 80% people in the world during their lifetime at some point, especially for the elderly^1-3^. However, it has shown a trend of affecting younger patients recently.^4^ The pathogenesis of the multi-factor condition remains unknown yet. The clinical therapeutic options for LBP mainly include conservative treatments and surgical operations, but none of them are satisfactory. As a result, it’s essential to explore the pathogenesis and underlying mechanisms of LBP for developing effective therapies.

LBP is mainly correlated with intervertebral disk degeneration and endplate injury^5,6^. In clinical, LBP caused by endplate injuries is quite common. Epidemiological investigations showed that, in the population with LBP, the incidence rate of endplate injures is about 30%^7^. During the growth and development of the vertebral body, the ossification of the epiphyseal plate on the upper and lower surfaces of the vertebral body stops, and the bone plate come into being, that is, the bony endplate. The center of the vertebral endplate is still covered by a thin layer of hyaline cartilage, which is the cartilage endplate^8,9^. In clinical MRI examination, the size of Modic changes and defects of cartilaginous and bony endplate are also strongly related with LBP^10-12^. It’s reported that, endplates change into porous during intervertebral disk degeneration. Recently, researchers found that, there are more nerve innervation and osteoclasts activation in cartilaginous endplate^13-15^. However, the dynamic condition in bony endplates were ignored. Drugs which could suppress the activities of osteoclasts such denosumab have shown analgesic effects in patients with Modic changes related LBP^14^. Therefore, the inhibition of osteoclastogenesis might prevent the local pathologic change and decrease the sensory innervation in endplates, which could be a potential strategy for LBP treatment^16^.

Growth differentiation factor 15 (GDF15) is a member of the transforming growth factor-β family that has been demonstrated to play an important role in several pathological state, such as cancer, inflammation, cardiovascular disease et al^17-20^. GDF15 is usually low expression under normal conditions in most tissues, but significantly increased in pathological conditions such as tissue injury or inflammation^21-23^. However, the effects of GDF15 on bone metabolism are still unclear. One study showed that, hypoxia was able to induce the expression of GDF15 and activated the NF-κB pathway, Mwhich promoted the osteoclasts differentiation^24^. In addition, GDF15 was also found to enhance osteoclastogenesis in human peripheral blood monocytes and tumor progression^25,26^. However, little is known on how GDF15 affects osteoclast differentiation in subchondral bone of cartilage endplate during low back pain.

Hence, in this study, we demonstrated that the LSI induced the overactivation of osteoclasts and CD31^hi^Emcn^hi^ endothelial vessels in the endplate. In addition, the expression of GDF15 in human disc samples and mice models were also increased. GDF15 could promote the fusion of preosteoclasts via Rac1/Cdc42/PAK/Cofilin axis, and facilitate the angiogenesis via the secretion of PDGF-BB. Furthermore, we proved that the GDF15 inhibitor, CTL-002 could reduce the expression of GDF15 in the endplate and alleviate the overactivation of osteoclasts and CD31^hi^Emcn^hi^ endothelial vessels induced by LSI in vivo. In conclusion, we demonstrated that GDF15 could regulates the fusion of preosteoclasts and targeting GDF15 in the endplate served as a novel anabolic therapy for low back pain treatment.

## Results

### 1. Construction of LBP originating from endplate models in LSI mice

We performed a lumbar spine instability (LSI) surgery to establish a mouse spine degeneration model for low back pain research. The behavior testing was examined at 2, 4 and 8 weeks after LSI surgery. We firstly explored the progression of low back pressure hyperalgesia. Compared with the sham group, the pressure tolerance decreased significantly after LSI surgery (Fig 1A). Then, we monitored the distance traveled, maximum speed and mean speed of movement per 24 h. The result showed that these spontaneous activities decreased significantly at 2, 4 and 8 weeks after LSI surgery (Fig 1B-1D). Furthermore, we examined the mechanical hyperalgesia of the hind paw by von Frey analysis as reported previously, which could indirectly reflect the severity of LBP. The paw withdraw frequency (PWF) in LSI group was significantly higher at 2, 4 and 8 weeks after LSI surgery (Fig 1E-1F). Above all, these results demonstrated that the spine instability model induces the development of LBP successfully.

**Figure 1.**
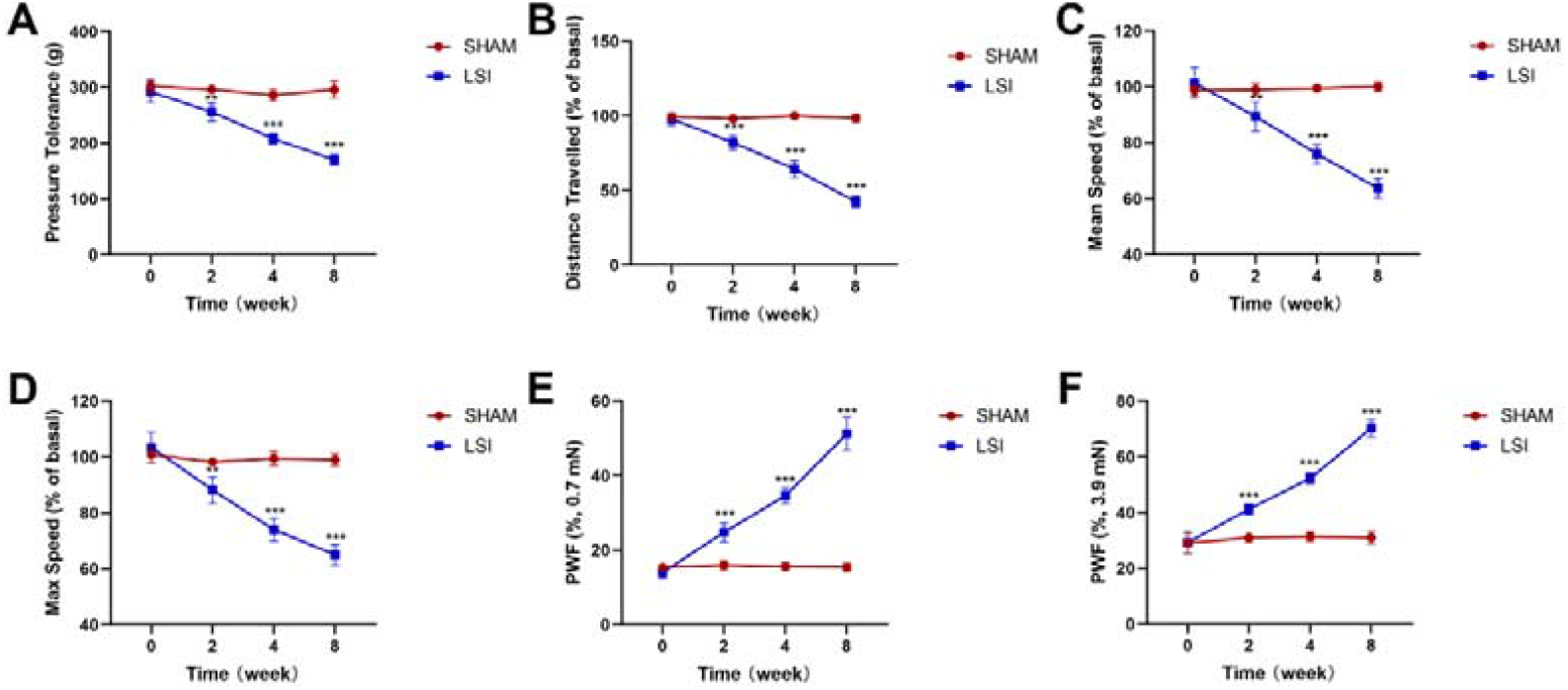
Symptomatic spinal pain behavior after the LSI surgery. **A**. The pressure hyperalgesia of the lumbar spine at different time point after sham or LSI surgery. The Spontaneous activity at different time points was evaluated by **B**. Distance traveled. **C**. Mean speed. **D**. Maximum speed. **E, F**. The hind paw withdrawal frequency responding to mechanical stimulation (von Frey, 0.7 mN and 3.9 mN) after sham or LSI surgery. n = 5 per group. (^*^P<0.05, ^**^P<0.01, ^***^P<0.001).

### 2. Sensory innervation in the endplate in LSI models

We then investigated the L4-L5 caudal endplate porosity in LSI models by micro-CT imaging and histological staining. The micro-CT analysis showed that, the micro-structure of the endplate in LSI group were more porous and the trabecular bone separation distribution (Tb. Sp) were higher compared with the sham group (Fig 2A-2C). For safranin O and fast green staining, the bone marrow surrounded the cavities of the endplates were stained green in LSI mice, indicating the ossification of the pathological endplate. The endplate scores were histologic assessment for pathological changes such as bone sclerosis, micro-structure disorders, which were significantly increased in LSI group (Fig 2D and 2E). The marker of peptidergic nociceptive C nerve fibers were detected by immunofluorescence staining, the results indicated that, the CGRP^+^ nerve fibers in the endplate were significantly increased at 2, 4 and 8 weeks after LSI surgery compared with the sham group (Fig 2F and 2G).

**Figure 2.**
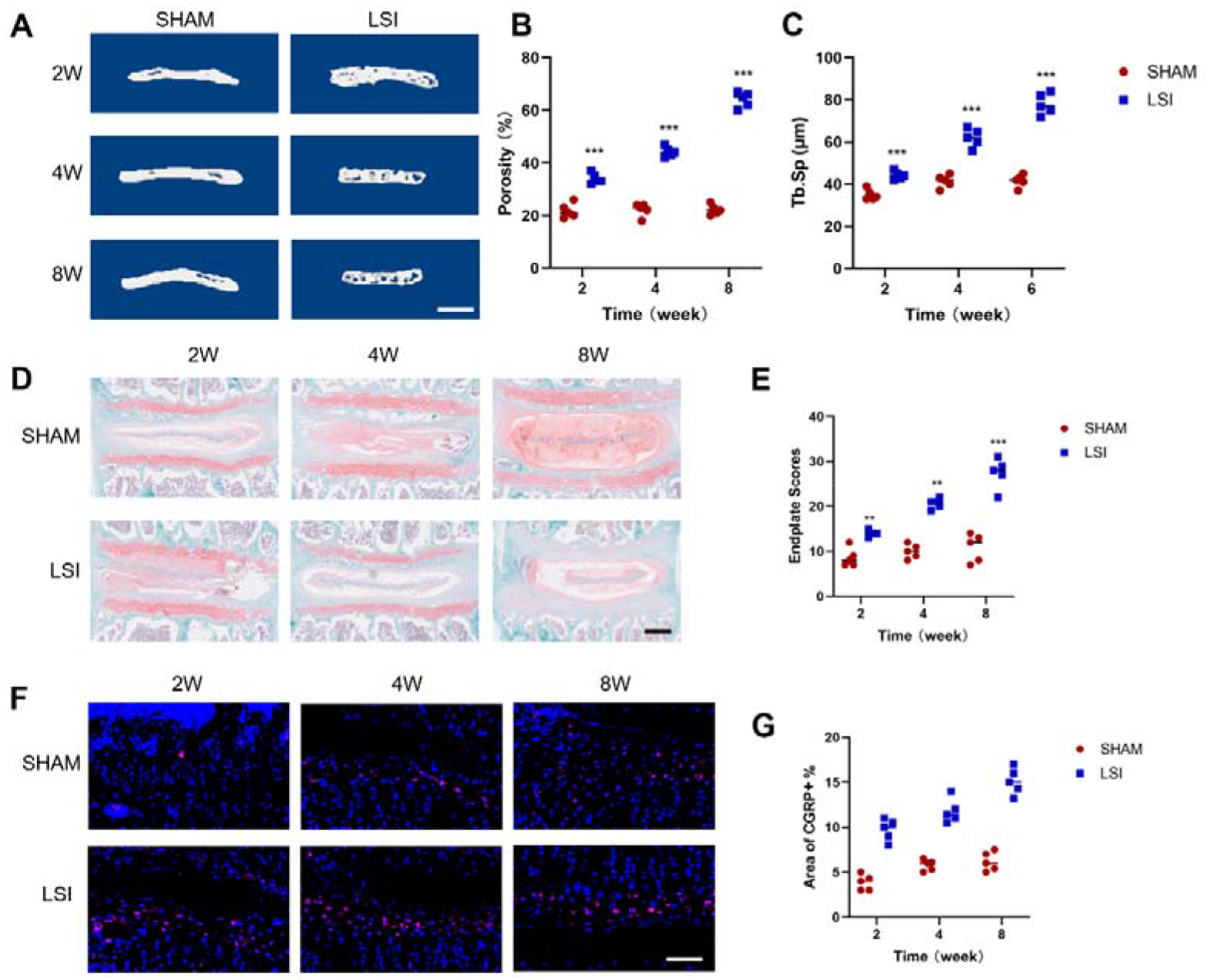
The CGRP^+^ nerve fibers were increased in endplates after LSI surgery. **A**. Representative three-dimensional micro-CT images of the mouse caudal endplates of L4/5 (coronal view) in sham or LSI mice. n=5 per group. Scale bars, 1mm. **B, C**. Quantitative analysis of the total porosity and trabecular separation (Tb. Sp) of the mouse caudal endplates of L4/5 determined by micro-CT. **D**. Representative images of safranin O and fast green staining of coronal caudal endplate sections of L4/5 in sham or LSI mice. n=5 per group. Scale bars, 200μm. **E**. Endplate scores based on safranin O and fast green staining. **F**. Representative images of CGRP^+^ nerve fibers (red) in mice caudal endplates of L4/5 at 2, 4 and 8 weeks after sham or LSI surgery. n=5 per group. Scale bars, 20μm. G. The percentage of CGRP^+^ nerve fiber area. (^*^P<0.05, ^**^P<0.01, ^***^P<0.001).

### 3. The osteoclatogensis and vascularization were over-activated in the bony endplate of LSI models

Previous study showed that the osteoclasts increased before endplate sclerosis^27^. To explore the bone metabolism changes in the endplate after LSI surgery. We examined the serum levels of osteogenesis and osteoclastogenesis markers. The results revealed the osteogenesis index of OCN levels after LSI surgery showed no significant change, however, the osteoclastogenesis index of Tracp 5b and CTX-1 increased at 2, 4 and 8 weeks after LSI surgery (Fig 3A-3C). These results demonstrated that the osteoclasts rather than osteoblast played the important role in LSI pathological progression in the early stages. We further performed TRAP staining to investigate the expression changes of osteoclasts in the bony endplate, the unfused mononuclear TRAP^+^ cells and multinuclear TRAP^+^ cells were analyzed. The results showed that, the unfused mononuclear TRAP+ cells were more abundant in the bony endplate in LSI group relative to the sham group (Fig 3D-3F). The H type vessels played an important role in bone metabolism balance. We performed immunofluorescence staining and the results showed that, compared with the sham group, the LSI mice had more CD31^hi^Emcn^hi^ cells in the endplate area at 2, 4 and 8 weeks after LSI operation (Fig 3G).

**Figure 3.**
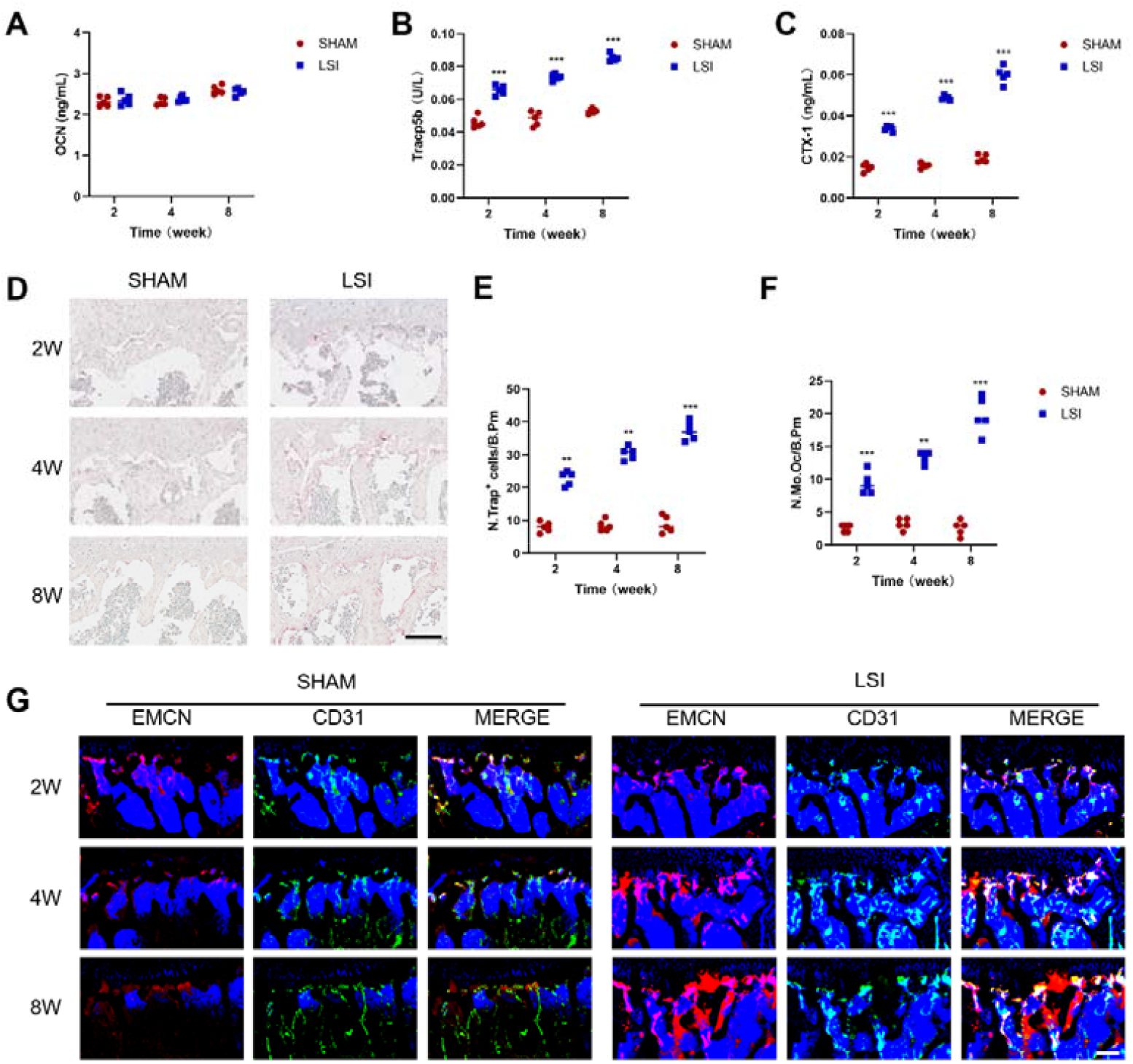
The osteoclasts are over-activated correlated with abundant H type vessels after LSI surgery. **A**. Serum levels of OCN. **B**. Serum levels of Tracp5b. **C**. Serum levels of CTX-1. **D**. Representative TRAP staining images of subchondral bone sections of L4/5 at 2, 4, and 8 weeks after LSI or sham surgery. n=5 per group. Scale bars, 50μm. **E, F**. Quantitative analysis of the number of TRAP^+^ mononuclear TRAP^+^ cells in endplates. **G**. Representative immunofluorescent images of endomucin (EMCN, red), CD31 (green) and EMCN^hi^ CD31^hi^ (yellow) cells of L4/5 at 2, 4, and 8 weeks after LSI or sham surgery. n=5 per group. Scale bars, 50μm. (^*^P<0.05, ^**^P<0.01, ^***^P<0.001).

### 4. The expression of GDF15 increased in LBP endplate tissues

LBP is mainly correlated with intervertebral disk degeneration-related diseases^5^. To explore the biological effects of different mRNAs in degenerative discs, we analyzed the microarray data (GSE70362) of 3 degenerative tissue samples versus 3 nondegenerative tissue samples. We pick out 114 significantly differentially expressed genes compared with the nondegenerative groups. We found that the growth differentiation factor 15 (GDF15) was one of the most significantly increased gene related to the differentiation of osteoclasts (Fig 4A-4C). In order to further confirmed the expression of GDF15 in LSI mice model, we performed immunohistochemical staining and the results revealed that, the GDF15 increased in the endplate at 2, 4 and 8 weeks after LSI surgery (Fig 4D).

**Figure 4.**
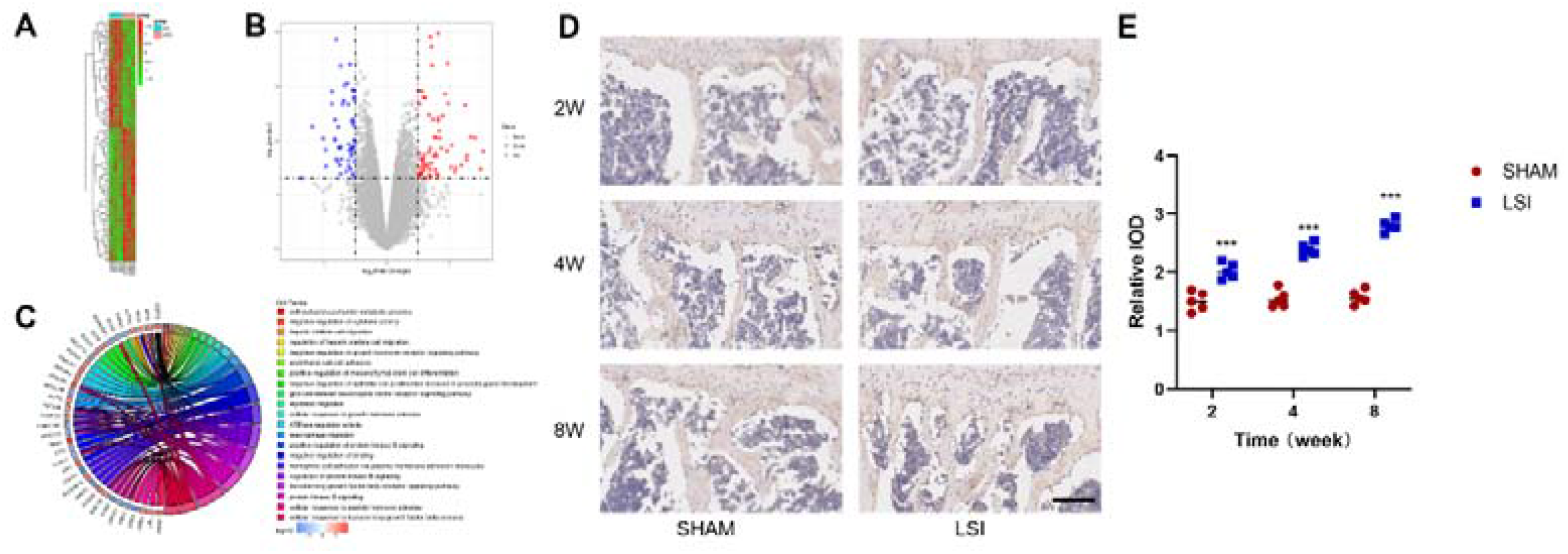
The expression of GDF15 was increased in low back pain patients and LSI mice. Identification of abnormally expressed genes in the GSE70362 dataset.**A**. Heatmap depicting all abnormally expressed genes in degenerative and nondegenerative intervertebral disc tissues. **B**. Volcano plot map of aberrantly expressed genes. **C**. Circos plot showing differentially expressed genes. **D**. The immunohistochemical staining of GDF15 at 2, 4 and 8 weeks after sham or LSI surgery. n=5 per group. Scale bars, 50μm. **E**. Quantitative analysis for the relative IOD of GDF15. (^*^P<0.05, ^**^P<0.01, ^***^P<0.001).

### 5. The GDF15 regulated the fusion of preosteoclasts by Rac1-Cdc42-PAK-Cofilin axis

As decreased multinuclear osteoclasts and increased preosteoclasts were detected in the endplate of LSI mice, and the expression of GDF15 increased after LSI surgery. We considered that the GDF15 played an important role during the fusion progression of osteoclasts. To test the conjecture, we separated the bone marrow monocytes and performed TRAP staining to detect the differentiation of osteoclasts at different time points. The results indicated that, the formation of preosteoclasts showed no statistical difference at day 3 after over-expressed GDF15 (Fig 5A and 5B). However, on day 6, there were more multinuclear osteoclasts in circ GDF15 group, indicating that the GDF15 is essential for the formation of mature osteoclasts rather than preosteoclasts (Fig 5C and 5D). We then performed cell fusion assay to investigate the role of GDF15 on the preosteoclasts fusion, cell membranes were labels by PKH67 (green) and PKH26 (red), 24 hours after staining, the fluorescence images were analyzed. The results showed that, GDF15 could promote the fusion of preosteoclasts obviously (Fig 5E and 5F). PDGF-BB was a key factor secreted by preosteoclasts, which played a significant prat in regulating angiogenesis and bone metabolism. We detected the cellular supernatant at different time points and found that, GDF15 could promote the secretion of GDF15 at day 3 and day 6 (Fig 5G). We then investigated the ability of angiogenesis by tube formation assay. The results indicated that conditioned medium from circ-GDF15 group induced more tube formation of endothelial progenitor cells (Fig 5H and 5I).

**Figure 5.**
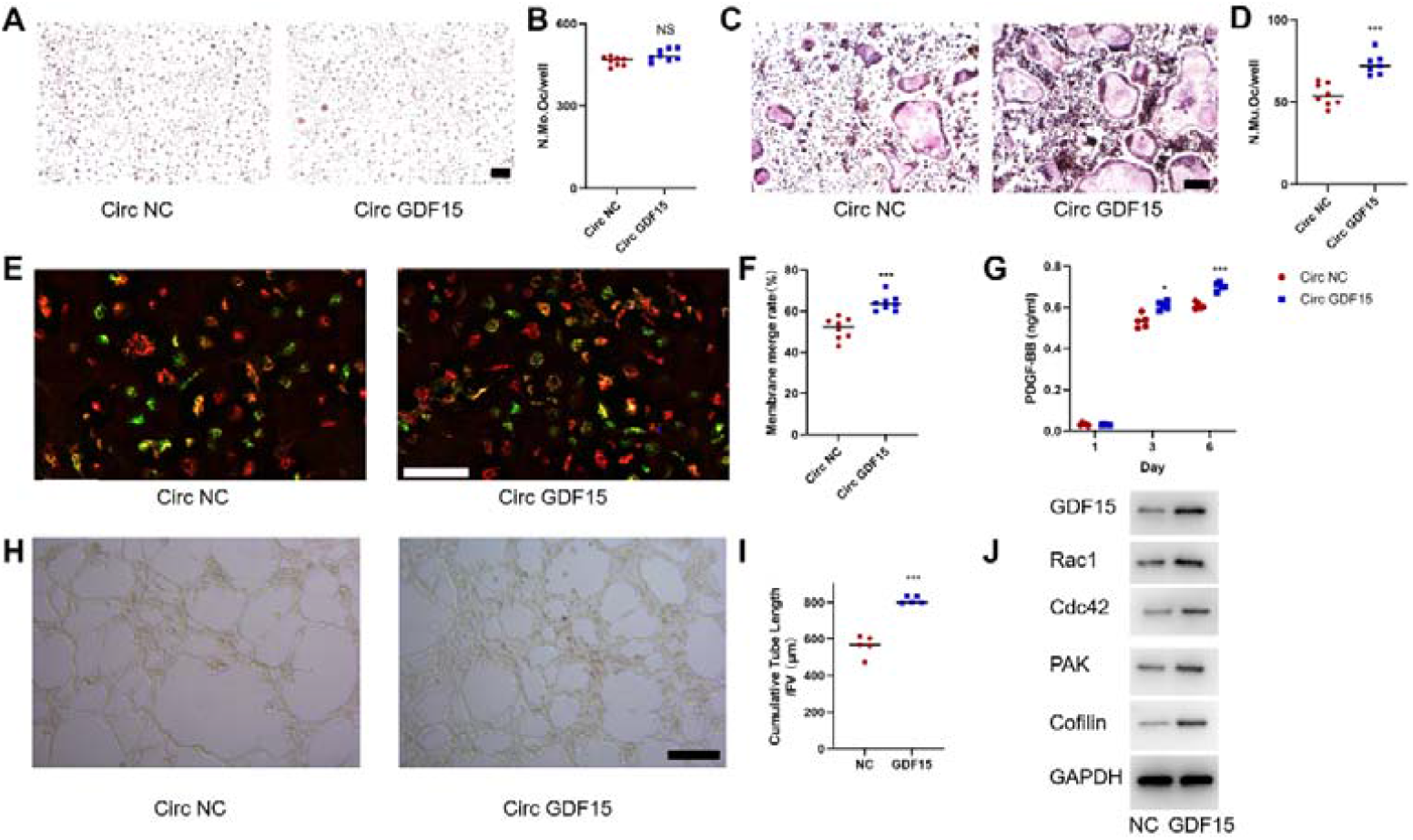
GDF15 is vital for preosteoclast fusion. **A**. TRAP staining of BMMs induced by RANKL on day 3. n=8 per group. Scale bar, 1000μm. **B**. Quantification of mononuclear TRAP^+^ cells. **C**. TRAP staining of BMMs induced by RANKL on day 6. n=8 per group. Scale bar, 200μm. **D**. Quantification of multinuclear TRAP^+^ cells. **E**. Representative images of double fluorescence for BMMs fusion on day 3. n=8 per group. Scale bar, 100μ m. **F**. Quantification of the membrane merge rate. **G**. Supernatant levels of PDGF-BB on days 1, 3 and 6 during osteoclast differentiation. n=5 per group. **H**. Matrigel tube formation assay with circ-NC or circ-GDF15 conditioned medium. n=5 per group. Scale bar, 200μm. **I**. Quantitative analysis of cumulative tube length. **J**. Western blot analysis for GDF15/Rac1/Cdc42/PAK/Cofilin of BMMs induced by RANKL on day 3. (^*^P<0.05, ^**^P<0.01, ^***^P<0.001).

The Rho GTP families played an important role in cell fusion, we then examined the expression of Rac1, Cdc42, PAK and Cofilin after GDF15 overexpression by western blot, the results showed that, after over-expression of GDF15, the Rho GTP enzymes were increased significantly (Fig 5J). We further investigated the role of rac1, cdc42 and PAK on the differentiation of osteoclasts. ZCL278 (cdc42 inhibitor), NSC23766 (rac1 inhibitor), FRAX597 (PAK inhibitor) were used to explore the cell fusion ability of preosteoclasts. The results showed that, ZCL278, NSC23766 and FRAX597 could reverse the effects of GDF15 overexpression on preosteoclasts fusion (Fig 6A and 6B). In addition, TRAP staining indicated that, ZCL278, NSC23766 and FRAX597 could reduce the formation of mature osteoclasts (Fig 6C and 6D). All these results revealed that, GDF15 promoted the fusion of preosteoclasts and osteoclastogenesis via rac1-cdc42-PAK-cofilin axis.

**Figure 6.**
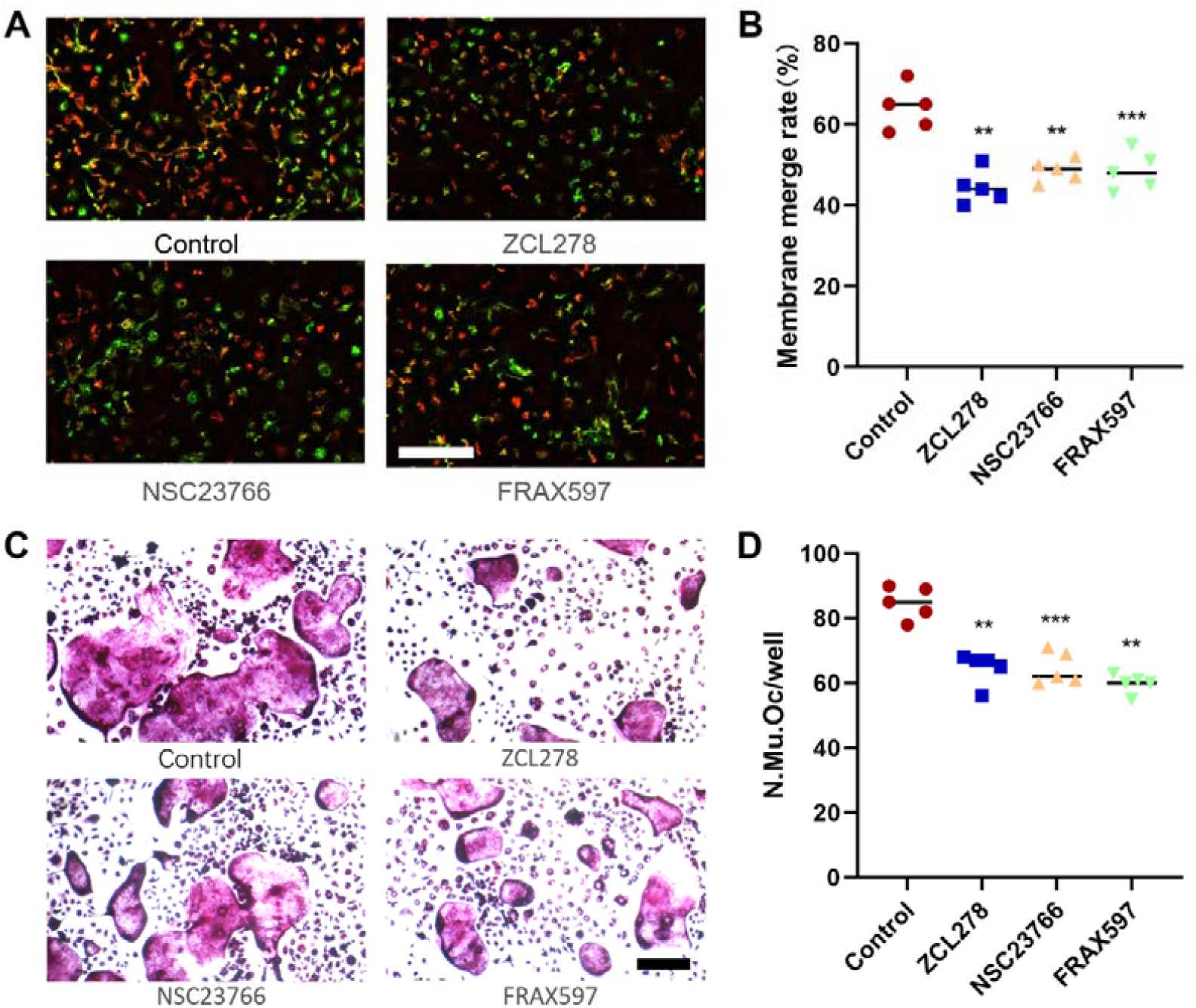
GDF15 is vital for preosteoclast fusion via Rac1/Cdc42/PAK/Cofilin axis. **A**. Representative images of double fluorescence for BMMs fusion on day 3. n=5 per group. Scale bar, 100μm. **B**. Quantification of the membrane merge rate. **C**. TRAP staining of BMMs induced by RANKL on day 6. n=5 per group. Scale bar, 200μm. **D**. Quantification of multinuclear TRAP^+^ cells. (^*^P<0.05, ^**^P<0.01, ^***^P<0.001).

### 6. Targeting GDF15 could suppress the over-activation of osteoclasts in the endplate and alleviate the low back pain originating from endplate

As GDF15 promoted the osteoclastogenesis and angiogenesis in vitro, we hypothesized that targeting GDF15 could be therapeutic potential for LBP treatment. CTL-002 was reported to be a GDF15 neutralizing antibody. In order to explore the role of GDF15 on LBP, we treated the mice with CTL-002 once a week for 8 weeks after LSI surgery. The immunochemistry staining showed that, the expression of GDF15 in the endplate decreased significantly after CTL-002 treatment (Fig 7A). The behavior testing was then examined at 2, 4 and 8 weeks after LSI surgery. We explored the progression of low back pressure hyperalgesia, compared with the control group, the pressure tolerance reversed significantly after CTL-002 treatment (Fig 7B). Then, we examined the distance traveled, maximum speed and mean speed of movement per 24 h. The result showed that these spontaneous activities reversed significantly at 2, 4 and 8 weeks after CTL-002 treatment (Fig 7C-7E). Furthermore, we examined the mechanical hyperalgesia of the hind paw. The PWF in LSI group was significantly higher at 2, 4 and 8 weeks after LSI surgery (Fig 7F and 7G). Above all, these results demonstrated that the spine instability induced by LSI was rescued by CTL-002.

**Figure 7.**
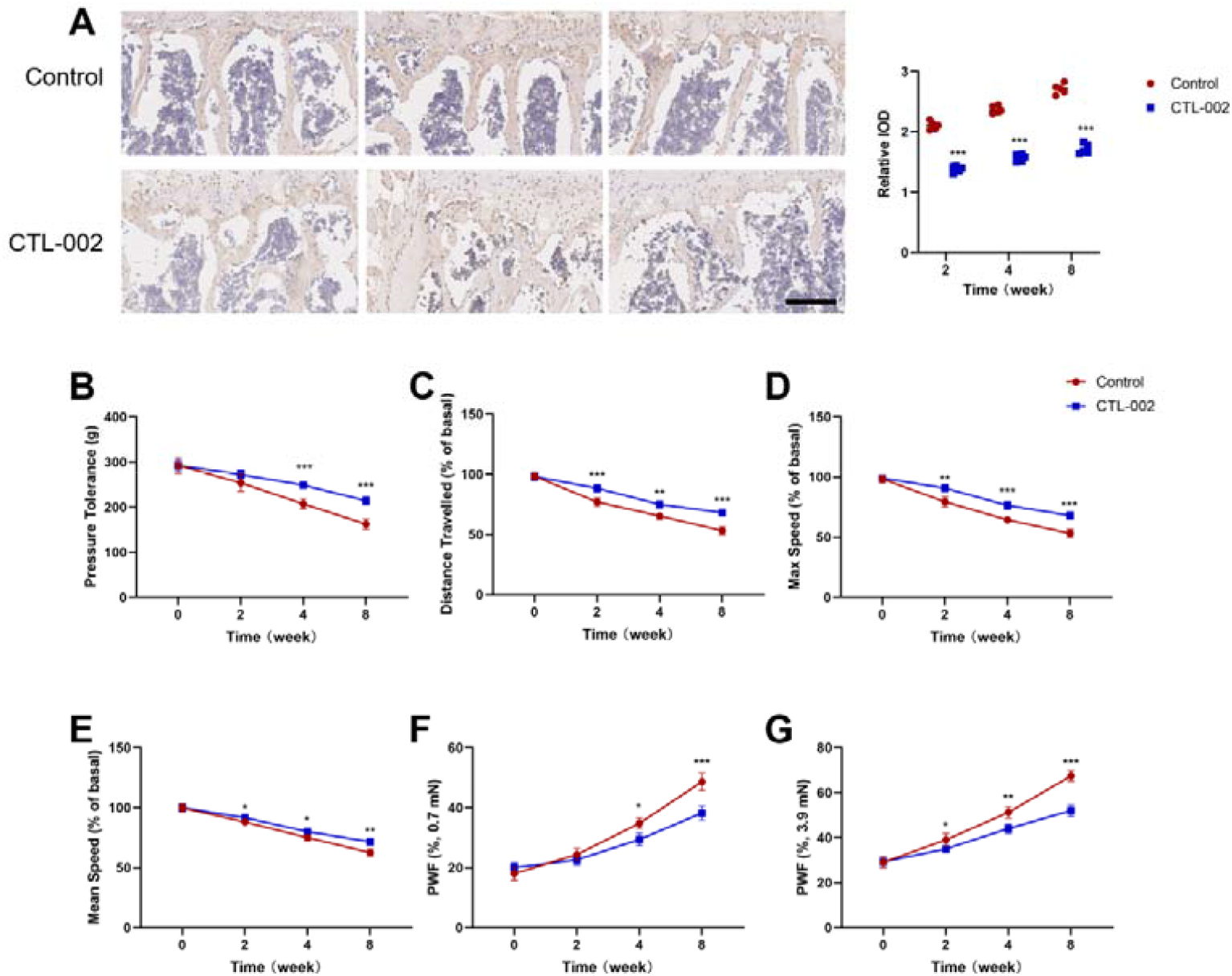
Targeting GDF15 could alleviate the symptomatic spinal pain behavior after the LSI surgery. **A**. The immunohistochemical staining of GDF15 at 2, 4 and 8 weeks after LSI surgery with control or CTL-002 treatment, and the quantitative analysis for the relative IOD of GDF15. n=5 per group. Scale bar, 200μm. **B**. The pressure hyperalgesia of the lumbar spine at different time point after control or CTL-002 treatment. The Spontaneous activity at different time points was evaluated by **C**. Distance traveled. **C**. Maximum speed. **D**. Mean speed. **E, F**. The hind paw withdrawal frequency responding to mechanical stimulation (von Frey, 0.7 mN and 3.9 mN) after control or CTL-002 treatment. n = 5 per group. (^*^P<0.05, ^**^P<0.01, ^***^P<0.001).

In order to investigate the osteoclastogenesis and angiogenesis in the endplate after targeting the GDF15, we performed TRAP and immunofluorescence staining. The results showed that, the unfused mononuclear TRAP^+^ cells were increased gradually at 2, 4 and 8 weeks after LSI surgery respectively. However, after targeting GDF15 by CTL-002, the number of mononuclear and multinuclear TRAP^+^ cells were reduced (Fig 8A-8C). In addition, the H type vessels in the endplate also decreased significantly compared with the control group, the mice in CTL-002 treatment group had less CD31^hi^Emcn^hi^ cells in the endplate area at 2, 4 and 8 weeks (Fig 8D).

**Figure 8.**
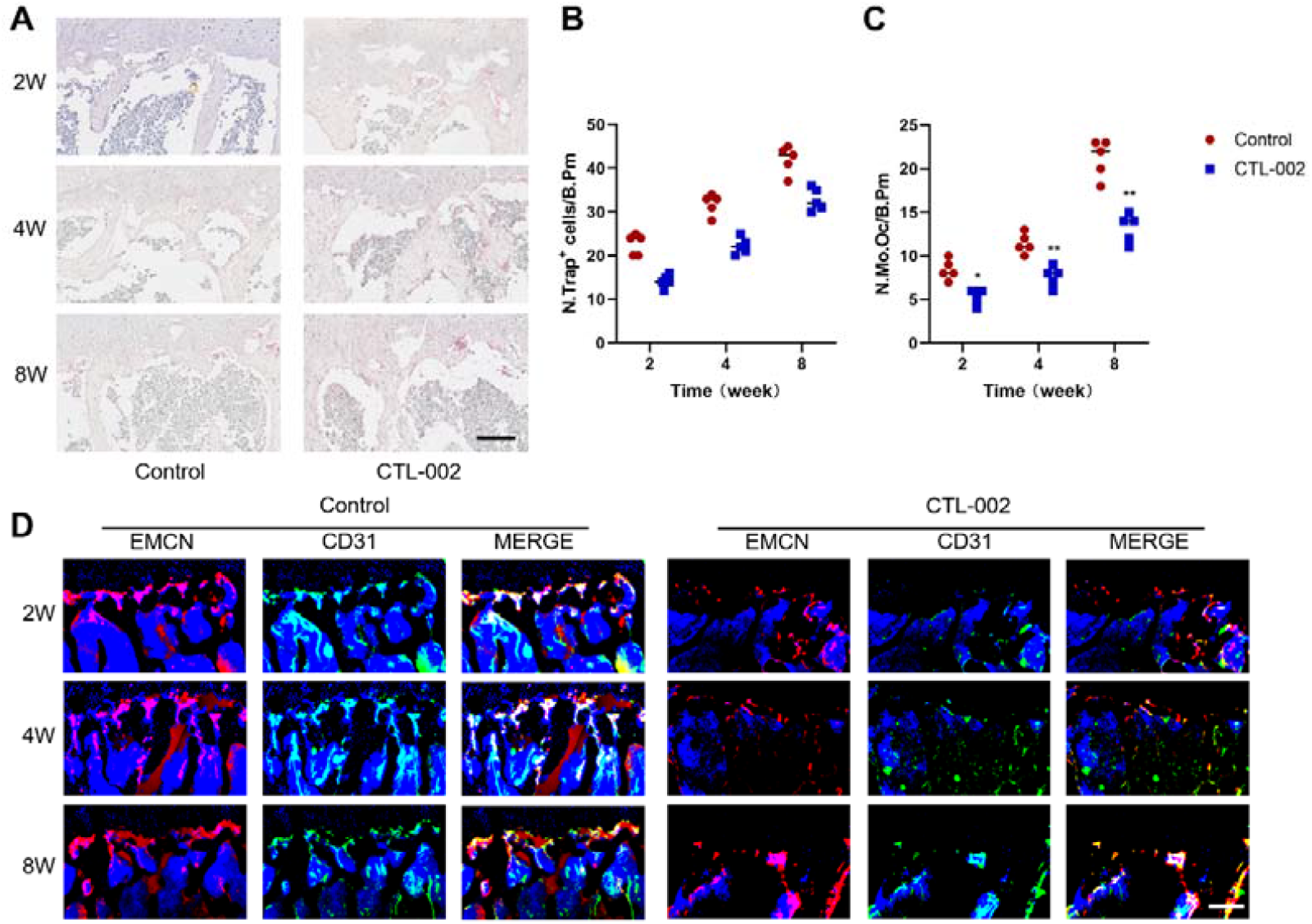
Targeting GDF15 could suppress the over-activation of osteoclasts. **A**. Representative TRAP staining images of subchondral bone sections of L4/5 at 2, 4, and 8 weeks after control or CTL-002 treatment. n=5 per group. Scale bars, 50μm. **B, C**. Quantitative analysis of the number of TRAP^+^ mononuclear TRAP^+^ cells in endplates. **D**. Representative immunofluorescent images of endomucin (EMCN, red), CD31 (green) and EMCN^hi^ CD31^hi^ (yellow) cells of L4/5 at 2, 4, and 8 weeks after control or CTL-002 treatment. n=5 per group. Scale bars, 50μm. (^*^P<0.05, ^**^P<0.01, ^***^P<0.001).

We then investigated the L4-L5 caudal endplate porosity after CTL-002 treatment by micro-CT analysis and histological staining. The micro-CT analysis showed that, the micro-structure of the endplate in CTL-002 treatment group were less porous and the trabecular bone separation distribution (Tb. Sp) were lower compared with the control group (Fig 9A-9C). For safranin O and fast green staining, the results showed that, the endplate scores significantly decreased after CTL-002 treatment (Fig 9D and 9E). The immunofluorescence staining of CGRP^+^ nerve fibers indicated that, the CGRP^+^ nerve fibers in the endplate were significantly increased at 2, 4 and 8 weeks after LSI surgery, however, after CTL-002 treatment, the CGRP^+^ area declined significantly (Fig 9F and 9G). In conclusion, targeting GDF15 could alleviate the over-activation of osteoclasts in the endplate and was supposed to be a potential therapy for the treatment of low back pain (Fig 10).

**Figure 9.**
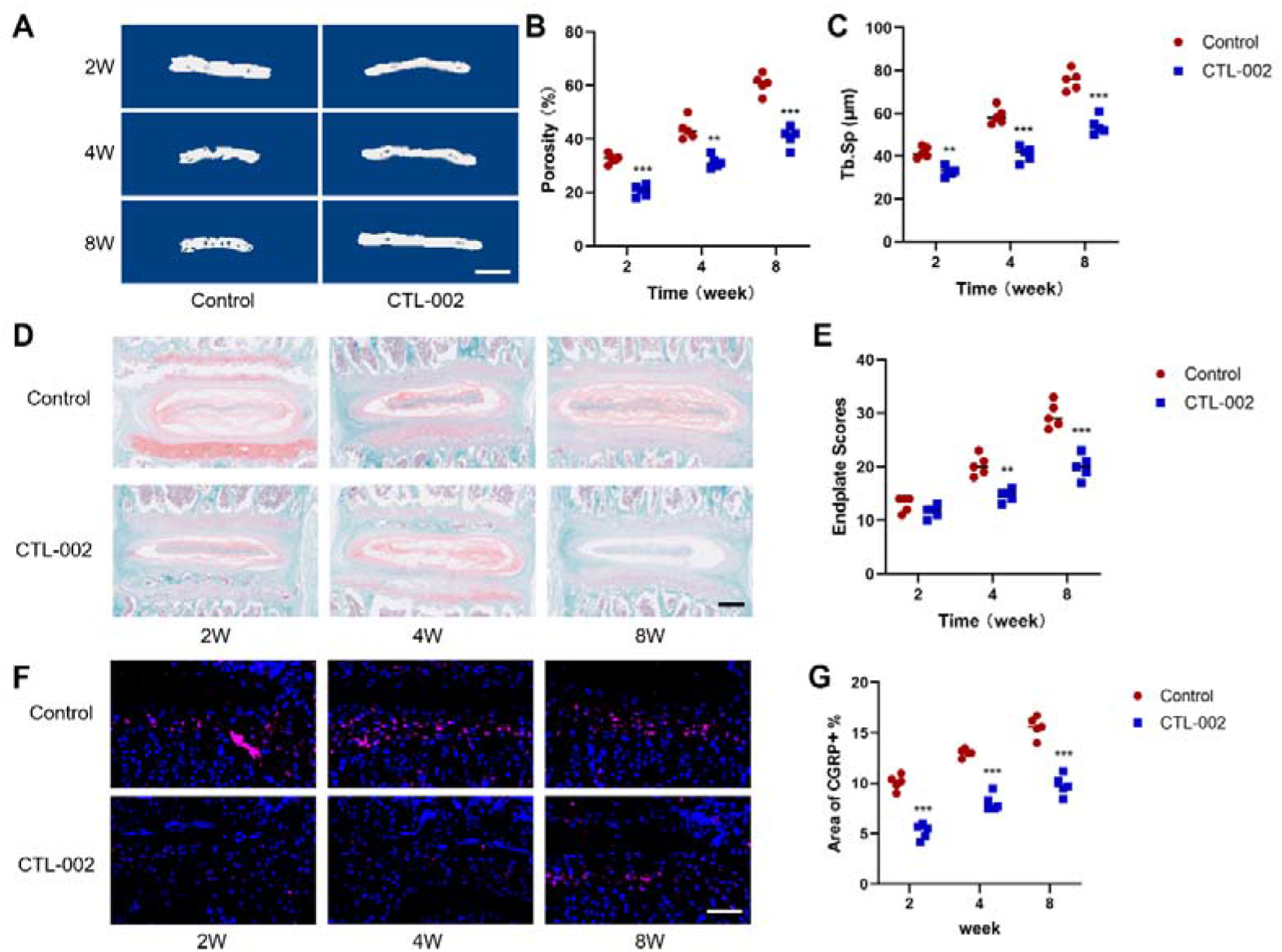
The sensory innervation in endplates decreased after targeting GDF15. **A**. Representative three-dimensional micro-CT images of the mouse caudal endplates of L4/5 (coronal view) in control or CTL-002 treatment. n=5 per group. Scale bars, 1mm. **B, C**. Quantitative analysis of the total porosity and trabecular separation (Tb. Sp) of the mouse caudal endplates of L4/5 determined by micro-CT. **D**. Representative images of safranin O and fast green staining of coronal caudal endplate sections of L4/5 in control or CTL-002 treatment. n=5 per group. Scale bars, 200μm. **E**. Endplate scores based on safranin O and fast green staining. **F**. Representative images of CGRP^+^ nerve fibers (red) in mice caudal endplates of L4/5 at 2, 4 and 8 weeks after control or CTL-002 treatment. n=5 per group. Scale bars, 20μm. **G**. The percentage of CGRP^+^ nerve fiber area. (^*^P<0.05, ^**^P<0.01, ^***^P<0.001).

**Figure 10.**
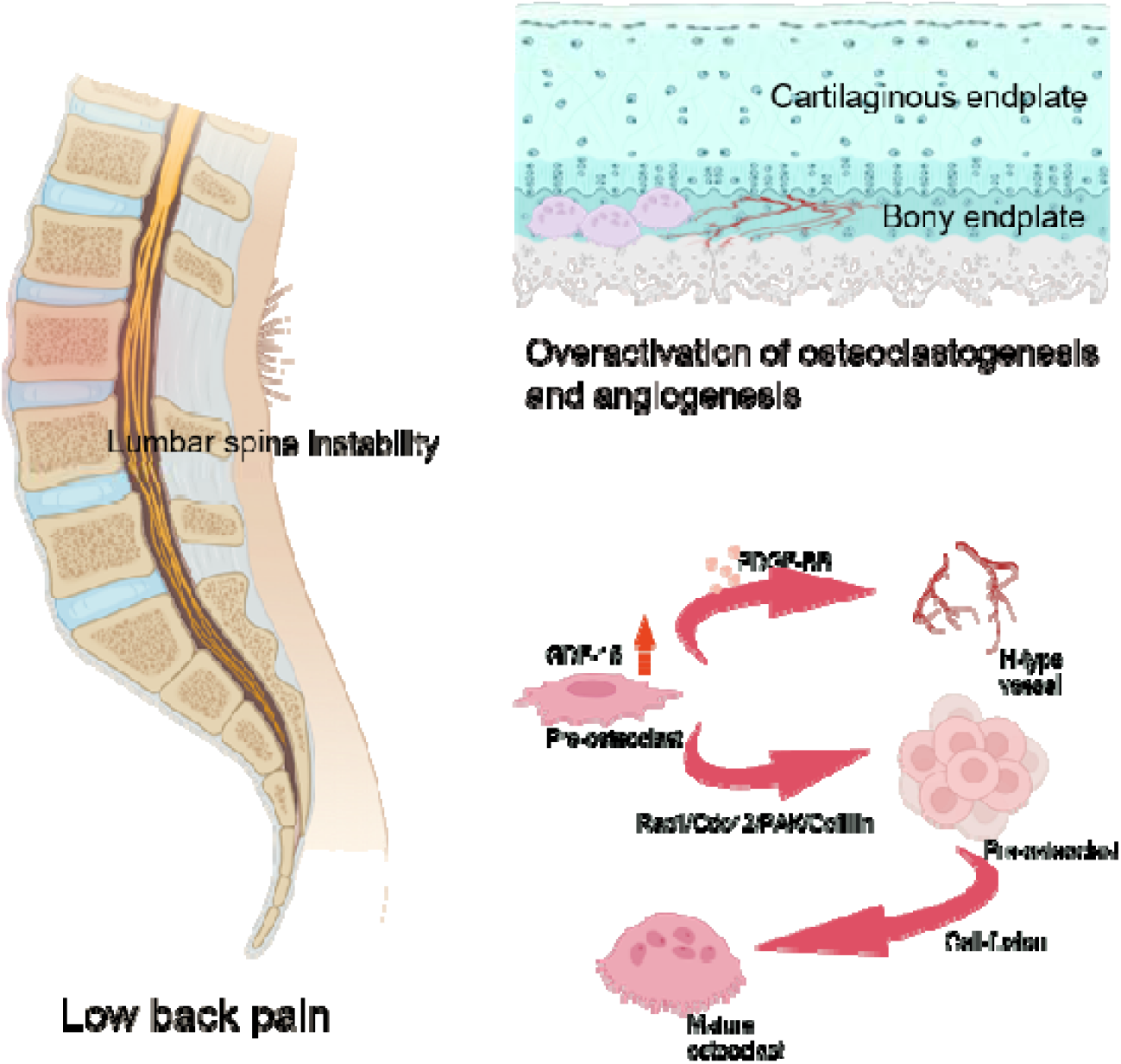
GDF15 facilitates osteoclast fusion through the Rac1/Cdc42/PAK/Cofilin pathway, targeting GDF15 as a potential therapy for low back pain originating from endplate.

## Materials

### Mouse models and in vivo experiments

8-week old C57BL/6 male mice were used for the in vivo experiments. In general, mice were firstly anesthetized by pentobarbital sodium (10mg/kg, intraperitoneally). Then, the LSI and sham models were conducted by excision of the L3-L5 spinous process, supraspinous and interspinous ligaments to result in the lumbar spine instability as reported previously. For the sham group, the operations were conducted by only detachment of the posterior paravertebral muscles from L3-L5 vertebral body.

### Micro-CT analyses

For quantitative micro-CT assays, mice were anesthetized with overdose of pentobarbital sodium. The lumbar spines were dissected and stored in 100% ethyl alcohol for 24 h and then stored in PBS solutions. All of the data were analyzed by a high resolution micro-CT (Skyscan 1076, Bruker). As reported previously ^28^, the scanner was set at a resolution of 8 m per pixel, a voltage of 80 kV, and a current of 124 A. The 3D models were analyzed by CTAn and CTVol. Two hundred section planes below the growth plate were observed.

### Immunofluorescence and histomorphometry

For immunofluorescence staining, the full lumbar spines were removed from the mice and fixed in 4% paraformalde-hyde at 4°C for 72 hours. Then, the spine tissues were decalcified with Ethylene Diamine Tetraacetic Acid (EDTA, 0.5 M; pH 7.4) solutions for 72 hours with continuous shaking. After that, the tissues were washed for at least 3 times with phosphate-buffered saline (PBS) solutions. Sections of 30 μm in the thickness were conducted for immunofluorescence staining. All of the experiments were conducted following with the previous studies^28,29^. Rat anti-EMCN antibody (1:100; sc-65495, Santa Cruz Biotechnology) and rabbit anti-CD31 antibody (1:20; ab28364, Abcam) were used for CD31 and EMCN immunofluorescence staining. Sections of 3 μm in the thickness were conducted for Safranin O and fast green and TRAP (Sigma-Aldrich) staining. Sections of 30 μm in the thickness were conducted for the CGRP sensory nerve immunofluorescent staining (1:100, ab81887, Abcam). All of the experiments were conducted following with the previous studies.^30^

### Western blotting

The primary antibodies for Western blotting analyses were rabbit anti-GDF15 (1:1000; ab206414, Abcam), rabbit anti–RAC1 (1:500; ab155938, Abcam), rabbit anti-Cdc42 (1:5000; ab187643, Abcam), rabbit anti–PAK (1:500; ab223849, Abcam), rabbit anti-Cofilin (1:2000; ab283500, Abcam). The secondary antibody was horseradish peroxidase–conjugated goat anti-rabbit (1:5000; ab6721, Abcam).

### Enzyme linked immunosorbent assay (ELISA)

The serum levels of CTX-1, TRACP-5b, OCN, PDGF-BB were detected by ELISA. The CTX-1 ELISA kit (NBP2-69074, Novus), TRACP-5b ELISA kit (EK7661, SAB), OCN ELISA kit (NBP2-68151, Novus) and PDGF-BB ELISA kit (ab224879, Abcam) were used respectively. All of the experiments were conducted following with the manufacturer’s instructions.

### Cell culture and transfection

The cells were incubated at 37°C in a humidified atmosphere with 5% CO_2_. Transfection was performed by circ-negative control, circ-GDF15. The forward oligonucleotide sequence is TAATACGACTCACTATAGGGCCACCATGCCCGGGCAAGAACTCAG; and the reverse oligonucleotide is TCTGAGATGAGTTTTTGTTCTATGCAGTGGCAGTCTTTGGCT.

### Osteoclast differentiation experiments

The bone marrow stromal cells were flushed out from the femurs of 8 weeks old C57BL/6 mice as described previously^31^. Then, cells were cultured for 3 days in α-MEM medium with 15% fetal bovine serum and 30 ng/mL M-CSF. Adherent cells were then collected and seeded into a 24 well plates. Cells were then incubated with M-CSF (30 ng/mL) and RANKL (100 ng/mL) for 3 (pre-osteoclast) and 6 (osteoclast) days respectively. Then, TRAP staining (Sigma-Aldrich, St. Louis, MO, USA) was conducted following the manufacturer’s instructions. TRAP-positive cells with more than 3 nuclei were regarded as mature osteoclasts.

### Osteoclastic fusion assays

The pre-osteoclasts were obtained as described above. The pre-osteoclasts were labeled with the red fluorescent dye PKH26 (Sigma, Aldrich) or the green florescent dye PKH67 (Sigma, Aldrich) following with the manufacturer’s instructions. Then, the two groups of pre-osteoclasts were seeded together on the 24 well plate for 2 hours. The liquid supernatant were removed and the fluorescence microscopy was conducted. The membrane merge rate was evaluated using the Image J as reported previously^32^.

### Tube formation experiments

The bone marrow stromal cells were flushed out from the femurs of 8 weeks old C57BL/6 mice as described previously. Then, cells were cultured for 3 days in α-MEM medium with 15% fetal bovine serum and 30 ng/mL M-CSF. Adherent cells were then collected and seeded into a 24 well plates. Cells were then incubated with M-CSF (30 ng/mL) and RANKL (100 ng/mL) for 3 (pre-osteoclast) days, and serum-containing conditioned medium was obtained. We put Matrigel (BD356231) in the plate and then polymerized for 3 hours. The EPCs (1 × 10^5^ cells per well) were seeded on polymerized Matrigel and followed with incubation of conditioned medium for 5 hours. After that, cells were stained by Calcein AM (C2012-0.1 ml, Beyotime) and the tube formation were analyzed by a fluorescent microscope.

### Behavioral Testing

All behavioral tests were conducted by the same investigator, who was blinded to the research groups.

Pressure tolerance was detected by the vocalization thresholds (as a nociceptive threshold) using a force gauge as described previously^30^. C57BL/6 mice were gently restrained and received the pressure force by a sensor on their skin above the L4-L5 spine. A cumulative increase of pressure force (50g/s) was conducted on the mice until the animals showed an audible vocalization.

Spontaneous running activity was measured by several indicators (including distance traveled, mean speed and maximum speed) by the activity wheels. C57BL/6 mice were placed in the cages which are similar to the mice’s home cages, and the wheels of the device could be used by the mice from all directions. The software of the equipment could show the real-time information of the spontaneous activity of the mice.

The pain hypersensitivity in response to external stimulation was detected by hind paw withdrawal frequency (PWF) using the von Frey test with 0.7 mN and 3.9 mN. C57BL/6 mice were put in a transparent plastic cage, which was put on a metal mesh grid. The midplantar position of the mice’s hind paw was stimulated by 0.7 mN or 3.9 mN. The frequency of mechanical stimulus was 10 times at a 1s interval. When the hind paw was withdrawn after the stimulation by von Frey filaments, the data was taken down.

### Statistical Analysis

We performed data analyses by using Graphpad Prism software. Data were presented as means ± SD or SEM, We used unpaired two-sample t-test to compare the data from 2 groups. We performed the one-way ANOVA to compare the data from multiple groups. P <0:05 was regarded as the statistical significance for all experiments.

## Discussion

LBP is a serious spondyloarthritis that is the leading cause of disability and is associated with high costs^33,34^. Existing treatments mainly included activity modification, operative treatment, and pharmaceutical agents to relieve the patient’s pain, but not to slow the LBP progression^35,36^. For operative treatment, such as total disc replacement and vertebral fusion, are the most effective clinical therapy. Most researchers paid attention to explore the pathogenesis of LBP on sensory innervation in the degenerative of lumbar intervertebral disc^37^. Nevertheless, the lumbar intervertebral disc degeneration is usually not accompanied by symptoms^38,39^. During the growth and development of the vertebral body, the ossification of the epiphyseal plate on the upper and lower surfaces of the vertebral body stops, and the bone plate come into being the bony endplate. The center of the vertebral endplate is still covered by a thin layer of hyaline cartilage, which is called the cartilage endplate^8,9^. Previous studies showed that, cartilaginous endplates was hardened and porous after up the aging process, which is clinically related to LBP^30^. Previous study showed that, the size of Modic changes and defects of cartilaginous and bony endplate were also strongly related with LBP^10-12^. The endplates changed into porous during intervertebral disk degeneration. Recently, researchers found that, there are more nerve innervation and osteoclasts activation in porous endplates of cartilaginous endplate^13-15^. Some studies demonstrated that the osteoclasts were overactivated up the aberrant mechanical loading process of the cartilage endplates^40,41^. It’s reported that, in the early stages of arthritis, the pre-osteoclasts were over-activated, and the osteoclastogenesis and angiogenesis were activated in the subchondral bone. However, the dynamic condition in bony endplates up the process of LSI were neglected. CD31^hi^Emcn^hi^ vessels were proved to be highly relevant to angiogenesis and osteogenesis. An increase of CD31^hi^Emcn^hi^ vessels were observed in the early stages of arthritis, and inhibition of CD31^hi^Emcn^hi^ vessels formation in the subchondral bone alleviated symptoms of arthritis^42,43^. The correlation between CD31^hi^Emcn^hi^ vessels and LBP were still not clarified. Monocytes could differentiate and merge into multinuclear osteoclasts under the M-CSF and RANKL culture. Prior to fusion into multinuclear osteoclasts, pre-osteoclasts are mononuclear TRAP positive cells, which produced PDGF-BB to bring CD31^hi^Emcn^hi^ vessels into being^28^. In this study, we firstly demonstrated that the osteoclasts and CD31^hi^Emcn^hi^ vessels were more abundant in bony endplate in LSI mice models, which is correlated with the pathological process of arthritis.

Growth differentiation factor 15 (GDF15) was identified from activated macrophage cell line clones, which act as an autocrine regulatory molecule in macrophages. It is a member of transforming growth factor β (TGFβ) superfamily. GDF15 molecules have been given different names according to their tissue origin or function at the time of discovery: macrophage inhibitory factor-1 (MIC-1), placental bone morphogenetic protein (PLAB), placental transforming growth factor-β (PTGF-β), non-steroidal anti-inflammatory drug activating gene-1 (NAG-1), prostate-derived growth factor (PDF). GDF15 is mainly involved in organ growth, differentiation, development and cell repair. Under physiological conditions, GDF15 is low expression in all tissues except the placenta. However, under pathological conditions such as inflammation or traumatic stress, GDF15 is up-regulated in the presence of TGFβ, interleukin 1β (IL-1β), tumor necrosis factor-α (TNF-α) and other stimulating factors et al^44,45^. A large number of studies have shown that elevated GDF15 levels are related to cardiovascular diseases such as myocardial hypertrophy, heart failure, atherosclerosis, endothelial dysfunction, as well as obesity, diabetes, cancer, cachexia, etc. GDF15 has been proved to be a new biomarker for the diagnosis, progression or prognosis of the above diseases and become a new target for the development of drugs^17-20,46,47^. Previous study showed that, GDF15 is a myokine in bone metabolism and muscle function in females with osteoporosis^23,48^. However, the role of GDF15 on LBP has not been demonstrated. In this study, we firstly reported that, the GDF15 is up-regulated in LBP patients and LSI mice models. Overexpression of GDF15 could promote the fusion of osteoclasts and secretion of PDGF-BB. CTL-002 is proved to be an effective neutralizing antibody of GDF15. In vivo studies showed that, CTL-002 treatment could reduce the expression of GDF15 in endplate, and targeting GDF15 could alleviate the overactivation of osteoclastogenesis and angiogenesis in bony endplate, and relieved the symptom of LBP significantly.

The dynamic polymerization of the cytoskeleton was actively involved in the migration, recognition and fusion of pre-osteoclasts, which played an important role in osteoclast differentiation^28,49,50^. Cytoskeleton was an important component of cell pseudopodia, and could be divided into lamellipodia and filopodia. The formation of cytoskeleton is mainly dependent on actin polymerization^51^. Thus, abnormalities in actin would lead to the chaos of cytoskeleton remodeling and affected cell migration and signal transduction, which eventually contributed to the abnormal of osteoclast differentiation. Rho GTPases signaling has been proved to be the key regulatory protein on actin polymerization^52,53^. Previous studies showed that, Rho GTPases played an important role on inflammation, tumor invasion, neurodegenerative disease and osteoclast differentiation. Cdc42 was one of the most important members of the Rho family and played an essential role in regulating the polymerization and depolymerization of cytoskeletal proteins^54,55^. Studies have shown that Cdc42 could regulate the cell migration, proliferation, differentiation and apoptosis via cell adhesion, cytoskeletal formation and reorganization. During the process of osteoclast differentiation, the cdc42 could promote the synthesis of F-actin and tubulin, which led to the protein polymerization and filopodia formation^56^. Rac and Cdc42 were the most specific Rho GTPases in the regulation of cell migration^57^. Rac could promote the assembly of lamellipodia. Cdc42 activated Rac and induced the recruitment of peripheral actin microfilaments, suggesting that there is a “crosstalk” between Rho GTPases^58^. During cell migration, the activation of Rac promoted the formation of actin protrusions at the leading edge of the cell and drove the actin contraction at the posterior edge of the cell. Then, Cdc42 regulates coordination with cell polarity in response to cues from extracellular orientation. Rac and Cdc42 had overlapping set of downstream effector molecules, which determined the changes in cytoskeleton. Including PAK and cofilin^59,60^. In this study, we found that, GDF15 could activated the Rac1/Cdc42/PAK/Cofilin axis. In cell fusion experiments, when circ-GDF15 was used in combination with Cdc42 inhibitor ZCL278, the Rac 1 inhibitor NSC23766, the PAK inhibitor FRAX597, the effects of GDF15 overexpression on preosteoclasts fusion were reversed significantly. In addition, TRAP staining indicated that, ZCL278, NSC23766 and FRAX597 could reduce the formation of mature osteoclasts. All these results revealed that, GDF15 promoted the fusion of preosteoclasts and osteoclastogenesis via rac1-cdc42-PAK-cofilin axis. Together, these results suggest that targeting GDF15 has the potential to treat low back pain as anabolic therapy (Fig. 10).

## Acknowledgement

This work was supported by the Natural Science Foundation of China (82201748), and Shanghai Sailing program (21YF1459200).

## Notes

### Competing Interest Statement

The authors have declared no competing interest.

